# Interhemispheric functional connectivity in the primary motor cortex distinguishes between training on a physical and a virtual surgical simulator

**DOI:** 10.1101/2021.07.10.451831

**Authors:** Anirban Dutta, Anil Kamat, Basiel Makled, Jack Norfleet, Xavier Intes, Suvranu De

## Abstract

Functional brain connectivity using functional near-infrared spectroscopy (fNIRS) during a pattern cutting (PC) task was investigated in physical and virtual simulators.

14 right-handed novice medical students were recruited and divided into separate cohorts for physical (N=8) and virtual (N=6) PC training. Functional brain connectivity measured were based on wavelet coherence (WCOH) from task-related oxygenated hemoglobin (HBO2) changes from baseline at left and right prefrontal cortex (LPFC, RPFC), left and right primary motor cortex (LPMC, RPMC), and supplementary motor area (SMA). HBO2 changes within the neurovascular frequency band (0.01-0.07Hz) from long-separation channels were used to compute average inter-regional WCOH metrics during the PC task. The coefficient of variation (CoV) of WCOH metrics and PC performance metrics were compared. WCOH metrics from short-separation fNIRS time-series were separately compared.

Partial eta squared effect size (Bonferroni correction) between the physical versus virtual simulator cohorts was found to be highest for LPMC-RPMC connectivity. Also, the percent change in magnitude-squared WCOH metric was statistically (p<0.05) different for LPMC-RPMC connectivity between the physical and the virtual simulator cohorts. Percent change in WCOH metrics from extracerebral sources was not different at the 5% significance level. Also, higher CoV for both LPMC-RPMC magnitude-squared WCOH metric and PC performance metrics were found in physical than a virtual simulator.

We conclude that interhemispheric connectivity of the primary motor cortex is the distinguishing functional brain connectivity feature between the physical versus the virtual simulator cohorts. Brain-behavior relationship based on CoV between the LPMC-RPMC magnitude-squared WCOH metric and the FLS PC performance metric provided novel insights into the neuroergonomics of the physical and virtual simulators that is crucial for validating Virtual Reality technology.

## 1 Introduction

Fundamentals of Laparoscopic Surgery (FLS) is a pre-requisite for board certification in general surgery in the USA. As a part of the FLS program, five psychomotor tasks with increasing task complexity are used for training and evaluation: (i) pegboard transfers, (ii) pattern cutting, (iii) placement of a ligating loop, (iv) suturing with extra-corporeal knot tying, and (v) suturing with intracorporeal knot tying. An important aspect of learning laparoscopic surgery compared to open surgery entails the sensorimotor and visuomotor adaptation of the 3D surgical field to 2D visualization, reduced tactile perception, and bimanual hand-eye coordination using ergonomically altered surgical instruments [1]. Prior works have shown the regional extent of brain activation in novices during laparoscopic surgery training in the functional magnetic resonance imaging (MRI) environment that depended on the task complexity [1] where tasks ii-v have shown activation of the Brodmann area (BA) 5 and BA 7 of the parietal cortex related to visual-motor coordination besides expected activation of BA 4 for bimanual coordination that includes the primary motor cortex (M1). While mainly the lateral portion of BA 6 of the frontal cortex showed activation for tasks i-iii, both the lateral and the medial portions of BA 6, including the premotor cortex and the supplementary motor area (SMA), showed activation in the most complex bimanually coordinated task v. In order to investigate these neural correlates (brain aspect) of laparoscopic surgery training, functional near infrared spectroscopy (fNIRS) can provide a better temporal resolution and ease of administration when compared to fMRI; however, at a lower spatial resolution and signal-to-noise ratio (SNR) [2]. Many prior works [3–7] have assessed surgery training using fNIRS mainly to compare skill levels; however, we could not find prior works on portable neuroimaging to compare surgery training in physical versus virtual simulators. This is crucial since surgical training field has seen a rapid emergence of new virtual training technologies which are gradually replacing physical training simulators with virtual training simulators. Here, virtual simulators been shown to improve acquisition of skills [8]; although, the neural correlates of skill acquisition in physical versus virtual simulators are unknown. Motor skill acquisition, consolidation, and long-term retention involve functional and structural plasticity where the temporal evolution of skill acquisition (behavioral aspect) can initially have a relatively fast phase followed by a later slower phase when further improvements develop incrementally over multiple sessions of practice [9].

In this study, we investigated fNIRS to capture brain activation during the pattern cutting (PC) task which may be considered a transitional task from the purely unimanual task i to the completely bimanual task v in terms of task complexity. In the PC task, the trainees cut a marked gauze and are given a score based on the accuracy of cutting and the time spent completing the activity. During the PC activity, the trainee provides traction to the gauze with one hand to place it in the best possible orientation to the cutting hand, and uses endoscopic scissors in the other hand to cut into the gauze along the pre-marked circle until it is completely removed. Therefore, modulation of interhemispheric inhibition of the M1 for bimanual coordination is expected during the PC task. Our prior work [10] investigated the brain functional connectivity based on wave-let coherence metrics for the following brain regions in right-handed subjects: left lateral prefrontal cortex, medial prefrontal cortex, right lateral prefrontal cortex, left medial primary motor cortex, and SMA, and found only in the case of FLS box trainer (not for virtual basic laparoscopic skills trainer: VBLaST) a correspondence to the surgical motor skills. Specifically, the partial eta squared effect size for the inter-regional magnitude-squared wavelet coherence metric between the medial prefrontal cortex and the SMA was found to be highest for expert versus novice comparison in the FLS box trainer cohort. Indeed, posterior SMA and right dorsal premotor area have been shown to be related to the bimanual coordination of finger movements [11] and medial prefrontal cortex is involved in long-term memory and decision making [12] expected to be different in experts versus novices. However, a lack of correspondence of brain functional connectivity to the surgical motor skills in VBLaST cohort needed further investigation since VBLaST has demonstrated face and construct validity [13, 14] including skill transfer [15]. Here, learning bimanual coordination of finger movements by novices for PC task performance (speed and accuracy) is postulated to involve learning to modulate interhemispheric inhibition of the primary motor cortices so we included fNIRS of bilateral primary motor cortices in the current study. Here, wavelet coherence [10] and cross-spectrum can provide a measure of the time-varying inter-regional association that may be sensitive to PC bimanual task-coupling to the interhemispheric coordination in physical versus virtual simulators. Therefore, we investigated fNIRS oxyhemoglobin (HbO2)-based wavelet coherence metric within the neurovascular frequency band (0.01-0.07Hz) related to neuronal activation, which was used to compare changes in the PC task-related brain functional connectivity from pre-task resting-state baseline in novices training in physical versus virtual simulators.

## 2 Materials and Methods

### 2.1 Subjects and experimental design

The study was approved by the Institutional Review Board of the Massachusetts General Hospital and the University at Buffalo. A priori power analysis, based on two-sample t-tests, determined the minimum number of samples required for this study. With a 95% confidence interval and a minimum power of 0.80, a minimum of eight subjects each for the expert and novice surgeon cohort group, four subjects for the FLS training group, three subjects for the VBLaST training group, and four subjects for the control group was estimated in our prior work [10]. In this study, we analyzed the data from 14 healthy novice medical students from prior work [10] where they performed performing the PC task in the physical and virtual simulators (demographics in the Table 1 of the Supplementary Materials). The novice medical students did not perform any laparoscopic procedures earlier. All right-handed subjects were divided into two cohorts; the physical simulator group consisted of 8 subjects whereas the virtual simulator group had 6 subjects. All the subjects were instructed verbally with a standard set of instructions on how to perform the PC task, goal, and the rule of the task completion. The optical probes or optodes held by a standard electroencephalography (EEG) cap (https://www.easycap.de) were mounted on the scalp of each participant by avoiding any hair in between the source/detector and the scalp. During the trial, the subjects were asked to perform the PC task, where the goal was to cut along the circular mark on a piece of gauze as accurately and as quickly as possible (details in our prior work [10]). After a baseline rest period of 1min, the trial was started and the maximum time provided for doing the PC task was 5 min. The scoring of the gauze cut on the physical simulator were subjectively assessed by the proctor, whereas for the virtual simulator the scores were automatically generated (details in our prior works [3, 10]).

### 2.2 Equipment

A 32-channel continuous-wave near-infrared spectrometer (CW6 system, TechEn Inc.) was used for this study that delivered infrared light at 690nm and 830nm. Optode montage consisted of eight long-distance and eight-short distance sources coupled to 16 detectors. 25 long-distance (30-40mm) channels and 8 short-distance (~8mm) channels measured brain activation at the bilateral prefrontal cortex (PFC), primary motor cortex (M1), and SMA that were assessed using Monte Carlo simulations in At-lasViewer (https://github.com/BUNPC/AtlasViewer). Based on the EEG cap locations and the optode sensitivity profile (Supplementary Materials – FigS1), we selected fNIRS channels that measured from the non-overlapping cortical regions (see Fig. 1), Left PFC (LPFC), Right PFC (RPFC), Left M1 (LPMC), Right M1 (RPMC), and SMA, based on anatomical guidance in AtlasViewer – an open-source software in Matlab (Mathworks Inc., USA) [9].

**Fig. 1.**
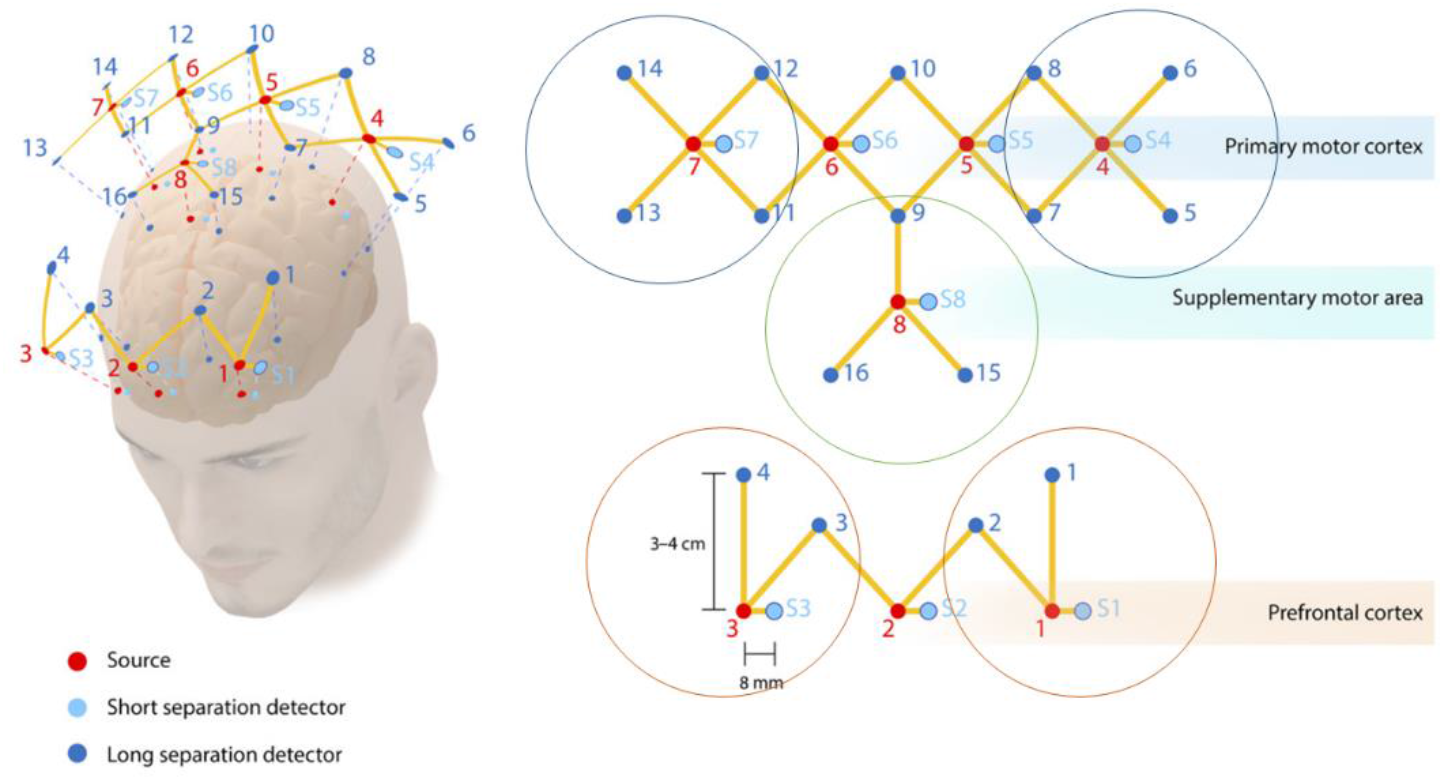
Selected optode positions over the PFC, M1, and SMA (from our prior work [3]) for nonoverlapping cortical regions, Left PFC (LPFC), Right PFC (RPFC), Left M1 (LPMC), Right M1 (RPMC), and SMA, based on anatomical guidance in AtlasViewer. Red dots indicate infrared sources, blue dots indicate long separation detectors, and light blue dots indicate short separation detectors. In the current study, PFC has two sources (1 and 3), two short separation detectors (S1 and S3), and four long separation detectors (1 to 4). The M1 has 2 sources (4 and 7), 2 short separation detectors (S4 and S7), and 8 long detectors (5 to 8 and 11 to 14). The SMA has one source (8), one short separation detector (S8), and three long separation detectors (9, 15, and 16).

### 2.3 Data processing for oxy-hemoglobin time-series

Motion artifact detection and correction were performed using combined spline interpolation and Savitzky-Golay filtering [10] in HOMER3 (https://github.com/BUNPC/Homer3), which is an open-source software in Matlab (Mathworks Inc., USA). Then, modified Beer-Lambert law was used to convert the detectors’ raw optical data into optical density. Then, the conversion of optical density to changes in HbO2 concentrations with differential path-length factors of 6.4 (690nm) and 5.8 (830nm) based on our prior work [3]. The mean HbO2 changes for each brain region [9], LPFC, RPFC, LPMC, RPMC, SMA, from long separation channels (inter-optode distance of 30-40mm) specific to the cortical activity in those regions was analyzed within neurovascular frequency band (0.01-0.07Hz) based on our prior work [10]. Short separation channels (inter-optode distance of 8mm) captured the systemic physiology originating from non-cortical superficial regions which was separately analyzed to investigate the effect of noise using the wavelet transform [16].

### 2.4 Mean and the coefficient of variation of functional connectivity metrics from fNIRS HbO2 time series and FLS performance scores

Pair-wise inter-regional functional connectivity between LPFC, RPFC, LPMC, RPMC, and SMA were estimated using wavelet coherence from the fNIRS HbO2 time series. In this study, wavelet coherence based pair-wise inter-regional functional connectivity was computed using the analytic Morlet wavelets that has shown good performance in time-frequency analysis by many prior works [10, 17, 18]. Analytic Morlet wavelets were centered within the neurovascular frequency band (0.01-0.07Hz) and applied using Wavelet Toolbox in Matlab (Mathworks Inc., USA). Due to the complex nature of the Morlet wavelet, the wavelet transform for each time and scale is a complex value. We performed the time-frequency analysis of the magnitude-squared wavelet coherence (WCOH), as shown in Fig. 2, and found the mean and the variance of WCOH along the time domain within the neurovascular frequency band (0.01-0.07Hz). Also, for a normalized measure of WCOH, the corresponding sine and cosine of the phase difference was averaged in time yielding wavelet phase coherence (WPCO) between the inter-regional HbO2 time series.

**Fig. 2.**
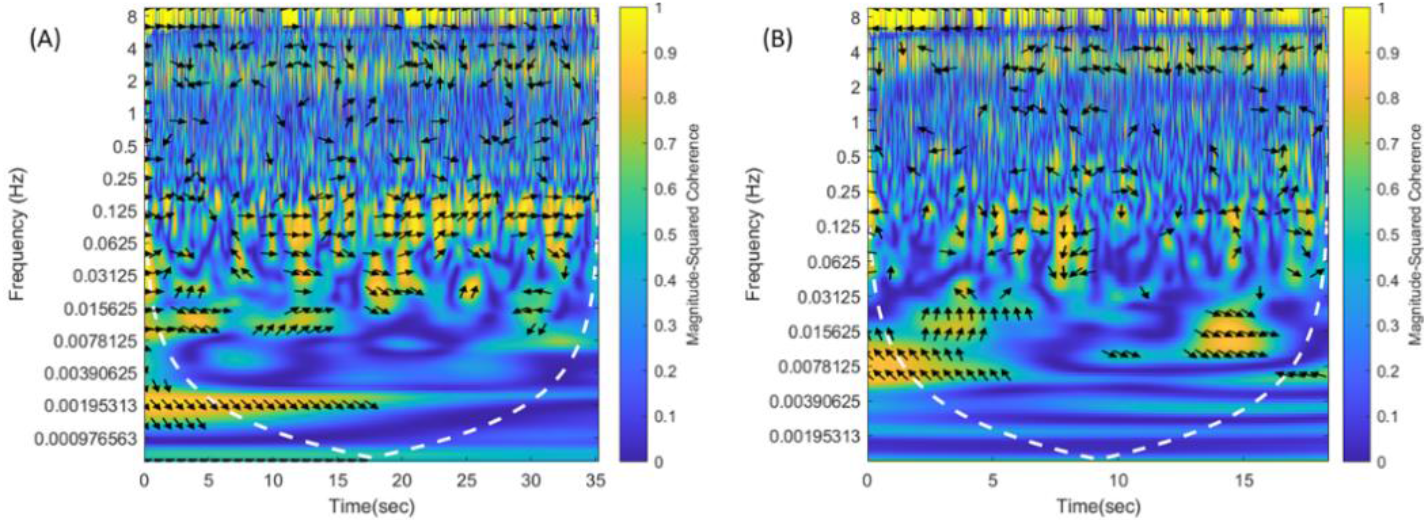
Illustrative examples of the magnitude-squared wavelet coherence between signals from the supplementary motor area and right primary cortex in the physical simulator (A) and virtual simulator (B). The color bar represents the squared magnitude.

The mean and the variance (also, coefficient of variation) of WCOH and WPCO were computed for the baseline resting state (1 min) and the PC task state separately from the long-separation and the short separation fNIRS channels. Here, brain-behavior relationship can be elucidated based on their coefficient of variation (CoV) [19], i.e., a higher CoV in the brain metrics (WCOH and WPCO) is postulated to be related to a higher CoV in the behavior metrics (FLS performance scores). Here, mean WCOH and WPCO values from the long-separation channels at the baseline and during the PC task estimated the functional connectivity for the following inter-regions: LPFC-RPFC, LPFC-LPMC, LPFC-RPMC, LPFC-SMA, RPFC-LPMC, RPFC-RPMC, RPFC-SMA, LPMC-RPMC, LPMC-SMA, RPMC-SMA.

The pre-task pair-wise mean WCOH and WPCO values were considered as the baseline connectivity and the percentage change from baseline was computed for the PC task state separately for the long-separation and the short-separation fNIRS channels. As a control measure, the percentage change from baseline during the PC task in the mean WCOH and WPCO values from the short separation channels were computed to determine systemic noise effects. After checks for normality (Shapiro–Wilk test, quantile-quantile plot) and homogeneity of variance (Levene test), we performed one-way multivariate analysis of variance (one-way MANOVA) in SPSS version 27 (IBM) to determine whether there is any significant difference in the inter-regional (i.e., LPFC-RPFC, LPFC-LPMC, LPFC-RPMC, LPFC-SMA, RPFC-LPMC, RPFC-RPMC, RPFC-SMA, LPMC-RPMC, LPMC-SMA, RPMC-SMA) functional connectivity metrics (average WCOH and WPCO values) using Wilks’ Lambda between the physical and the virtual simulator cohorts. Then, to determine how the dependent variables (i.e., inter-regional functional connectivity) differ for the independent variable (physical versus virtual simulator), partial eta squared effect size was used with alpha correction with Bonferroni correction. Then, one-way analysis of variance (ANOVA) was performed to determine whether the independent variable, the physical (FLSbox) and the virtual (VBLaST) simulators have different effects on individual inter-regional functional connectivity (percent change in magnitude-squared WCOH metric) separately for the long-separation and the short-separation fNIRS channels.

## 3 Results

### 3.1 Physical versus virtual simulator effect on the functional connectivity

An illustrative example of the magnitude-squared wavelet coherence plot for RPMC-SMA in the physical and virtual simulators is shown in the Fig. 2. Grouped plot of the matrix of percentage change from baseline in average WCOH and WPCO during PC task in physical (FLSbox) and virtual (VBLaST) simulators across ten inter-regional LPFC-RPFC, LPFC-LPMC, LPFC-RPMC, LPFC-SMA, RPFC-LPMC, RPFC-RPMC, RPFC-SMA, LPMC-RPMC, LPMC-SMA, RPMC-SMA are shown in the Supplementary materials (FigS2). One-way multivariate ANOVA did not find significant differences at the 1% level in the averages of ten (inter-regional) WCOH and ten (inter-regional) WPCO measures between the physical (FLSbox) and the virtual (VBLaST) simulators. Partial eta squared effect size (Bonferroni correction) was found to be highest for LPMC-RPMC connectivity.

### 3.2 Statistical analysis on physical versus virtual simulator effect on the individual inter-regional functional connectivity

Shapiro-Wilk test of normality conducted on the functional brain connectivity metrics (WCOH, WPCO) found that while all the metrics in the virtual simulator cohort passed the test at the 5% level, few metrics in the physical simulator cohort digressed from the normal distribution. Therefore, we performed both the one-way ANOVA and the Wilcoxon Rank Sum Test to determine whether the independent variable, the physical (FLSbox) and the virtual (VBLaST) simulators, have different effects on the individual response variables, WCOH and WPCO measures. Results are summarized in the Fig. S3 (Supplementary Materials), where the percent change from baseline in the WCOH metric for LPMC – RPMC functional connectivity was found to be statistically (p < 0.05) different between the physical and the virtual simulators for the long separation channels. However, no significant differences at the 5% level in the percent change from baseline in the percent change from baseline in the WCOH metric were found for the short separation channels. Also, no significant differences at the 5% level in the percent change from baseline in the WPCO metrics between the physical and the virtual simulator cohorts were found for both the long and the short separation channels.

### 3.3 Brain – behavior correspondence in terms of the coefficient of variation

Fig. 3. shows that the physical (FLSbox) resulted in higher CoV than the virtual (VBLaST) simulator in both the brain (WCOH, WPCO) and the behavior metrics (FLS performance scores). Specifically, CoV for the WCOH measure (see Fig. 3B) for the LPMC-RPMC and the CoV for the WPCO measure for the RPMC-SMA (see Fig. 3C) were highest in the physical (FLSbox) simulator.

**Fig. 3.**
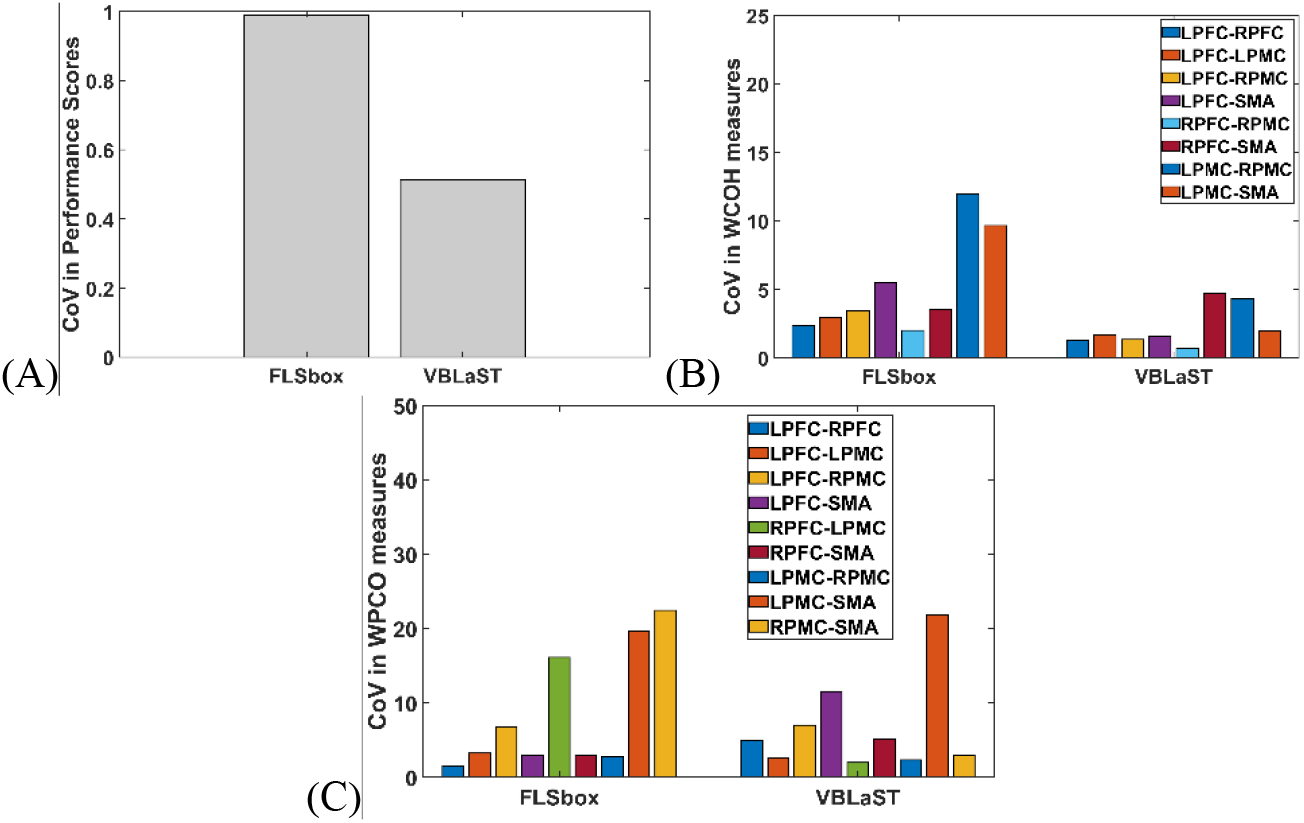
Coefficient of variation (CoV) in brain (WCOH, WPCO) and behavior (FLS performance) measures. [*WCOH, WPCO outliers removed based on ‘isoutlier’ in Matlab*] (A) CoV for the FLS performance scores in the physical (FLSbox) and the virtual (VBLaST) simulators. (B) CoV for the percent change from baseline in inter-regional magnitude-squared wavelet coherence (WCOH) measures in FLSbox and VBLaST simulators. (C) CoV for percent change from baseline in inter-regional wavelet phase coherence (WPCO) measures in FLSbox and VBLaST simulators.

## 4 Discussion

WCOH metric for the LPMC-RPMC functional connectivity was the best marker between FLS PC training in the physical and the virtual simulators in terms of both partial eta squared effect size and the CoV-based brain-behavior analysis (see Fig. 3). Importantly, higher CoV for the LPMC-RPMC functional connectivity in the physical simulator corresponded with higher CoV in FLS performance score in the physical simulator, as shown in Fig. 3. Even with our limitation due to a small sample size (that affected MANOVA results), our partial eta squared effect size and the CoV-based brain-behavior analysis highlighted the relevance of LPMC-RPMC functional connectivity during FLS PC training. Here, nomological validity can be derived from prior work [19] that showed cortical functional MRI variability in parietal cortex explained the movement extent variability. Also, the relevance of inter-hemispheric LPMC-RPMC functional connectivity is supported by the relevance of corpus callosum in bimanual coordination [20].

In our study, novice subjects performed more variable movements in the physical simulator than the virtual simulator that was related to higher variability in the functional connectivity metrics in the physical simulator. Our results also aligned well with the prior work using whole-brain imaging [1] that have demonstrated the necessity of the modulation of interhemispheric inhibition (primary motor cortices) for bimanual coordination. The crucial brain and behavior correspondence was found based on CoV [19], i.e., relative variability in interhemispheric LPMC-RPMC functional connectivity is postulated as the “neural correlate” of the relative variability in the FLS performance score. Based on current study results showing brain and behavior correspondence based on CoV between physical versus virtual simulator, it is postulated that in the early phase of skill acquisition, the CoV of PFC functional brain connectivity will decrease corresponding to a decrease in the CoV of FLS performance score with increasing familiarity [21]. Here, activation in the PFC is expected in the early phase of skill acquisition when attention and working memory are required to actively monitor targets in the environment until ‘automaticity’ is achieved.

In this study we have highlighted the need to study the variability in the brain and behavior metrics and their correspondence which is rarely studied [19]. Moreover, this brain and behavior relationship in terms of CoV was established based on fNIRS in this study when compared to functional MRI [19]. Although whole-brain imaging [1] has been shown feasible during FLS task, performing FLS task in a functional MRI environment required modification of the FLS environment, restricting whole-body mobility while also requiring an abnormal posture. Conversely, fNIRS provided a portable low-cost solution without interfering with the certified FLS task environment. This is also relevant for neuroimaging guided transcranial electrical stimulation (tES) [22] where we have shown the feasibility of online monitoring of the brain activation during surgical training that can be facilitated with tES [23–25][13], [14]. Such portable neuroimaging guided tES approach, based on the brain-behavior correspondence, will allow subject-specific learning-stage specific application using multi-channel tES [26]. Our brain – behavior approach based on the correspondence of CoV for brain and behavior metrics can be used to find neural correlates (brain aspect) to assess and improve laparoscopic surgical skills. Therefore, we postulate that fNIRS-based investigation of the variability in behavior is crucial in neuroergonomics for studying the brain and behavior at work [27]. For future studies on the brain-behavior relationship, dynamic modulation of the interhemispheric inhibition of primary motor cortices during motor learning of bimanual (alternating movement initiation and inhibition, e.g., in FLS suturing tasks) coordination also needs further investigation during more complex laparoscopic skill training.

## 5 Conclusion

We showed that inter-hemispheric primary motor cortex functional connectivity based on wavelet coherence was most dissimilar in novices between the PC task performed in the physical versus virtual simulators. Also, the brain-behavior relationship was found based on the CoV in the physical versus virtual simulator where the normalized or relative measure of the variation was found higher in the physical simulator than the virtual simulator.

## 6 Acknowledgment

The authors gratefully acknowledge the support of this work through the Medical Technology Enterprise Consortium (MTEC) award #W81XWH2090019 (2020-628), and the U.S. Army Futures Command, Combat Capabilities Development Command Soldier Center STTC cooperative research agreement #W912CG-21-2-0001. This work was also partly supported by the National Institute of Biomedical Imaging and Bioengineering (NIBIB) under Grant Nos. 1R01EB014305, NHLBI 1R01HL119248, and NCI 1R01CA197491 grants. We would also like to thank Arthur “Buzz” DiMartino and his team at TechEn for graciously providing us support with the CW6 spectrometer.

## References

1. Bahrami, P., Graham, S.J., Grantcharov, T.P., Cusimano, M.D., Rotstein, O.D., Mansur, A., Schweizer, T.A.: Neuroanatomical correlates of laparoscopic surgery training. Surg Endosc. 28, 2189–2198 (2014). https://doi.org/10.1007/s00464-014-3452-7.

2. Cui, X., Bray, S., Bryant, D.M., Glover, G.H., Reiss, A.L.: A quantitative comparison of NIRS and fMRI across multiple cognitive tasks. Neuroimage. 54, 2808–2821 (2011). https://doi.org/10.1016/j.neuroimage.2010.10.069.

3. Nemani, A., Yücel, M.A., Kruger, U., Gee, D.W., Cooper, C., Schwaitzberg, S.D., De, S., Intes, X.: Assessing bimanual motor skills with optical neuroimaging. Science Advances. 4, eaat3807 (2018). https://doi.org/10.1126/sciadv.aat3807.

4. Nemani, A., Kruger, U., Cooper, C.A., Schwaitzberg, S.D., Intes, X., De, S.: Objective assessment of surgical skill transfer using non-invasive brain imaging. Surg Endosc. 33, 2485–2494 (2019). https://doi.org/10.1007/s00464-018-6535-z.

5. Khoe, H.C.H., Low, J.W., Wijerathne, S., Ann, L.S., Salgaonkar, H., Lomanto, D., Choi, J., Baek, J., Tam, W.W., Pei, H., Ho, R.C.M.: Use of prefrontal cortex activity as a measure of learning curve in surgical novices: results of a single blind randomised controlled trial. Surg Endosc. 34, 5604–5615 (2020). https://doi.org/10.1007/s00464-019-07331-7.

6. Leff, D.R., Orihuela-Espina, F., Atallah, L., Athanasiou, T., Leong, J.J.H., Darzi, A.W., Yang, G.-Z.: Functional prefrontal reorganization accompanies learning-associated refinements in surgery: a manifold embedding approach. Comput Aided Surg. 13, 325–339 (2008). https://doi.org/10.3109/10929080802531482.

7. Keles, H.O., Cengiz, C., Demiral, I., Ozmen, M.M., Omurtag, A.: High density optical neuroimaging predicts surgeons’s subjective experience and skill levels. PLOS ONE. 16, e0247117 (2021). https://doi.org/10.1371/journal.pone.0247117.

8. Jordan, J.A., Gallagher, A.G., McGuigan, J., McClure, N.: Virtual reality training leads to faster adaptation to the novel psychomotor restrictions encountered by laparoscopic surgeons. Surg Endosc. 15, 1080–1084 (2001). https://doi.org/10.1007/s004640000374.

9. Dayan, E., Cohen, L.G.: Neuroplasticity subserving motor skill learning. Neuron. 72, 443–454 (2011). https://doi.org/10.1016/j.neuron.2011.10.008.

10. Nemani, A., Kamat, A., Gao, Y., Yucel, M.A., Gee, D., Cooper, C., Schwaitzberg, S.D., Intes, X., Dutta, A., De, S.: Functional brain connectivity related to surgical skill dexterity in physical and virtual simulation environments. NPh. 8, 015008 (2021). https://doi.org/10.1117/1.NPh.8.1.015008.

11. Sadato, N., Yonekura, Y., Waki, A., Yamada, H., Ishii, Y.: Role of the Supplementary Motor Area and the Right Premotor Cortex in the Coordination of Bimanual Finger Movements. J Neurosci. 17, 9667–9674 (1997). https://doi.org/10.1523/JNEUROSCI.17-24-09667.1997.

12. Euston, D.R., Gruber, A.J., McNaughton, B.L.: The Role of Medial Prefrontal Cortex in Memory and Decision Making. Neuron. 76, 1057–1070 (2012). https://doi.org/10.1016/j.neuron.2012.12.002.

13. Sankaranarayanan, G., Lin, H., Arikatla, V.S., Mulcare, M., Zhang, L., Derevianko, A., Lim, R., Fobert, D., Cao, C., Schwaitzberg, S.D., Jones, D.B., De, S.: Preliminary Face and Construct Validation Study of a Virtual Basic Laparoscopic Skill Trainer. J Laparoendosc Adv Surg Tech A. 20, 153–157 (2010). https://doi.org/10.1089/lap.2009.0030.

14. Linsk, A.M., Monden, K.R., Sankaranarayanan, G., Ahn, W., Jones, D.B., De, S., Schwaitzberg, S.D., Cao, C.G.L.: Validation of the VBLaST pattern cutting task: a learning curve study. Surg Endosc. 32, 1990–2002 (2018). https://doi.org/10.1007/s00464-017-5895-0.

15. Nemani, A., Ahn, W., Cooper, C., Schwaitzberg, S., De, S.: Convergent validation and transfer of learning studies of a virtual reality-based pattern cutting simulator. Surg Endosc. 32, 1265–1272 (2018). https://doi.org/10.1007/s00464-017-5802-8.

16. Duan, L., Zhao, Z., Lin, Y., Wu, X., Luo, Y., Xu, P.: Wavelet-based method for removing global physiological noise in functional near-infrared spectroscopy. Biomed. Opt. Express, BOE. 9, 3805–3820 (2018). https://doi.org/10.1364/BOE.9.003805.

17. Frontiers | Identifying and quantifying main components of physiological noise in functional near infrared spectroscopy on the prefrontal cortex | Human Neuroscience, https://www.frontiersin.org/articles/10.3389/fnhum.2013.00864/full, last accessed 2021/05/21.

18. Zhang, X., Yu, J., Zhao, R., Xu, W., Niu, H., Zhang, Y., Zuo, N., Jiang, T.: Activation detection in functional near-infrared spectroscopy by wavelet coherence. J Biomed Opt. 20, 016004 (2015). https://doi.org/10.1117/1.JBO.20.1.016004.

19. Haar, S., Donchin, O., Dinstein, I.: Individual Movement Variability Magnitudes Are Explained by Cortical Neural Variability. J. Neurosci. 37, 9076–9085 (2017). https://doi.org/10.1523/JNEUROSCI.1650-17.2017.

20. Gooijers, J., Swinnen, S.P.: Interactions between brain structure and behavior: the corpus callosum and bimanual coordination. Neurosci Biobehav Rev. 43, 1–19 (2014). https://doi.org/10.1016/j.neubiorev.2014.03.008.

21. Leff, D.R., Elwell, C.E., Orihuela-Espina, F., Atallah, L., Delpy, D.T., Darzi, A.W., Yang, G.Z.: Changes in prefrontal cortical behaviour depend upon familiarity on a bimanual co-ordination task: an fNIRS study. Neuroimage. 39, 805–813 (2008). https://doi.org/10.1016/j.neuroimage.2007.09.032.

22. Guhathakurta, D., Dutta, A.: Computational Pipeline for NIRS-EEG Joint Imaging of tDCS-Evoked Cerebral Responses—An Application in Ischemic Stroke. Front. Neurosci. 10, (2016). https://doi.org/10.3389/fnins.2016.00261.

23. Patel, R., Singh, H., Ashcroft, J., Woods, A.J., Darzi, A., Leff, D.R.: Dataset of Prefrontal Transcranial Direct-Current Stimulation to Improve Early Surgical Knot-tying Skills. Data in Brief. 106905 (2021). https://doi.org/10.1016/j.dib.2021.106905.

24. Ashcroft, J., Patel, R., Woods, A.J., Darzi, A., Singh, H., Leff, D.R.: Prefrontal transcranial direct-current stimulation improves early technical skills in surgery. Brain Stimul. 13, 1834–1841 (2020). https://doi.org/10.1016/j.brs.2020.10.013.

25. Gao, Y., Cavuoto, L., Schwaitzberg, S., Norfleet, J.E., Intes, X., De, S.: The Effects of Transcranial Electrical Stimulation on Human Motor Functions: A Comprehensive Review of Functional Neuroimaging Studies. Front Neurosci. 14, (2020). https://doi.org/10.3389/fnins.2020.00744.

26. Otal, B., Dutta, A., Foerster, Á., Ripolles, O., Kuceyeski, A., Miranda, P.C., Edwards, D.J., Ilić, T.V., Nitsche, M.A., Ruffini, G.: Opportunities for Guided Multichannel Non-invasive Transcranial Current Stimulation in Poststroke Rehabilitation. Front Neurol. 7, (2016). https://doi.org/10.3389/fneur.2016.00021.

27. Dehais, F., Lafont, A., Roy, R., Fairclough, S.: A Neuroergonomics Approach to Mental Workload, Engagement and Human Performance. Front Neurosci. 14, (2020). https://doi.org/10.3389/fnins.2020.00268.

